# Targeted delivery of galunisertib attenuates fibrogenesis in an integrated *ex vivo* renal transplant and fibrosis model

**DOI:** 10.1101/2022.03.22.485255

**Authors:** L. Leonie van Leeuwen, Henri G.D. Leuvenink, Benedikt M. Kessler, Peter Olinga, Mitchel J.R. Ruigrok

## Abstract

Normothermic machine perfusion is an emerging preservation technique for kidney allografts to reduce post-transplant complications, including interstitial fibrosis and tubular atrophy. This technique, however, could be improved by adding antifibrotic molecules to perfusion solutions. We established Machine perfusion and Organ slices as a Platform for Ex vivo Drug delivery (MOPED), to explore fibrogenesis suppression strategies. We perfused porcine kidneys *ex vivo* with galunisertib—a potent inhibitor of the transforming growth factor beta signaling pathway. To determine whether effects persisted, we also cultured precision-cut tissue slices prepared from the respective kidneys. Galunisertib supplementation improved the general viability, without negatively affecting renal function or elevating levels of injury markers or byproducts of oxidative stress. Galunisertib also reduced inflammation and more importantly, strongly suppressed the onset of fibrosis, especially when the treatment was continued in slices. Our results illustrate the value of targeted drug delivery, using isolated organ perfusion, for reducing post-transplant complications.

**One Sentence Summary:** Galunisertib supplementation during normothermic machine perfusion attenuates fibrogenesis without compromising renal function.

## INTRODUCTION

Kidney transplantation is a life-saving procedure for patients that suffer from end-stage renal disease, which is characterized by severe microalbuminuria and vastly reduced glomerular filtration *(1)*. Unfortunately, not all patients are eligible for a kidney transplant, as the demand far exceeds the supply *(2)*. On top of that, the clinical outcomes of kidney transplantation are not always good as post-transplant complications are frequently observed. One of the most common post-transplant complications is interstitial fibrosis and tubular atrophy (IF/TA). In fact, this complication is detectable in ~25% of kidney allografts after 1 year, and in 90% after 10 years *(3)*. Patients who develop IF/TA eventually have to resume dialysis, undergo re-transplantation, or suffer from premature death, as the allograft function declines due to the excessive deposition of extracellular matrix proteins and loss of tubular epithelial cells. Therefore, safer and more effective treatments for slowing—or perhaps even reversing— IF/TA are greatly desired.

One approach to reduce the onset of IF/TA is to minimize allograft damage caused by ischemia-reperfusion injury (IRI). To do so, clinicians are increasingly using machine perfusion, as it has been demonstrated to be superior to static cold storage *(4, 5)*. The concept of machine perfusion is based on the controlled flow of a solution, containing nutrients, metabolites, and oxygen through an *ex vivo* organ. Besides hypothermic machine perfusion (HMP) which aims to preserve organs, normothermic machine perfusion (NMP) technology has been introduced in the clinics to assess organ function *(6)*. This technique could also be used as a treatment platform by supplementing perfusion solutions with inhibitors of signaling pathways that regulate fibrogenesis, such as galunisertib, which is an inhibitor of the transforming growth factor beta (TGF-β) pathway *(2)*. The TGF-β pathway plays a key role in fibrosis because it drives the differentiation of fibroblasts into myofibroblasts—key effector cells that produce large quantities of matrix proteins, especially collagens and fibronectins *(7)*. For this reason, galunisertib seems to be a promising drug candidate for attenuating fibrosis during NMP.

We therefore investigated the safety and efficacy of this novel therapeutic approach, using a newly developed drug testing platform. We present Machine perfusion and Organ slices as a Platform for *Ex vivo* Drug delivery (MOPED), a robust technique for testing the efficacy of *ex vivo* (anti-fibrotic) therapies. We perfused porcine kidneys for 6 hours with a blood-based perfusate containing TGF-β1, galunisertib, or a combination thereof. To determine whether effects persisted upon ceasing or continuing treatments, we also cultured precision-cut tissue slices, prepared from the treated kidneys, for an additional 48 hours. Slices are viable explants that can be cultured *ex vivo* for up to a few days while retaining functional and structural characteristics, thereby offering an unmatched opportunity to further explore the potential effects of galunisertib on renal tissue *(8, 9)*. With respect to readouts, we focused on analyzing the general tissue viability and renal function, as well as the release of general injury markers and specific oxidative stress markers. We also comprehensively characterized the effect of galunisertib on the extent of inflammation and, more importantly, the onset of fibrogenesis.

## RESULTS

### Setup of an *ex vivo* renal transplant and fibrosis model

Our integrated *ex vivo* model, optimal for drug delivery, was based on state-of-the-art machine perfusion techniques combined with the use of precision-cut tissue slices, entitled MOPED (Fig. 1A). We obtained porcine kidneys, with an average weight of 365 ± 36 g, from a local abattoir and subjected them to 30 min of warm ischemia to mimic similar circumstances as during deceased donation. The kidneys were then transported and preserved for 24 h by means of oxygenated HMP, using a mean arterial pressure of 25 mmHg and a temperature of 3.8 ± 1.2 °C (Fig. 1B). During HMP, the flow rate ranged between 25-70 mL/min and increased over time. We subsequently performed NMP using a custom-built setup, configured with a mean arterial pressure of 80 mmHg and a temperature of 37 ± 1.3°C (Fig. 1C). Kidneys were subjected to 1 h of NMP before adding treatments to the perfusate (i.e., TGF-β1, galunisertib, or a combination thereof). TGF-β1 was added to the perfusate to induce fibrogenesis. During NMP, the flow rate ranged from 200 to 500 mL/min. The renal flow rate was significantly lower for kidneys perfused with TGF-β1 as compared to other groups. After NMP, the kidney tissue was used for the preparation of slices. This was performed to establish whether potential effects persisted upon ceasing or continuing treatments.

**FIG. 1.**
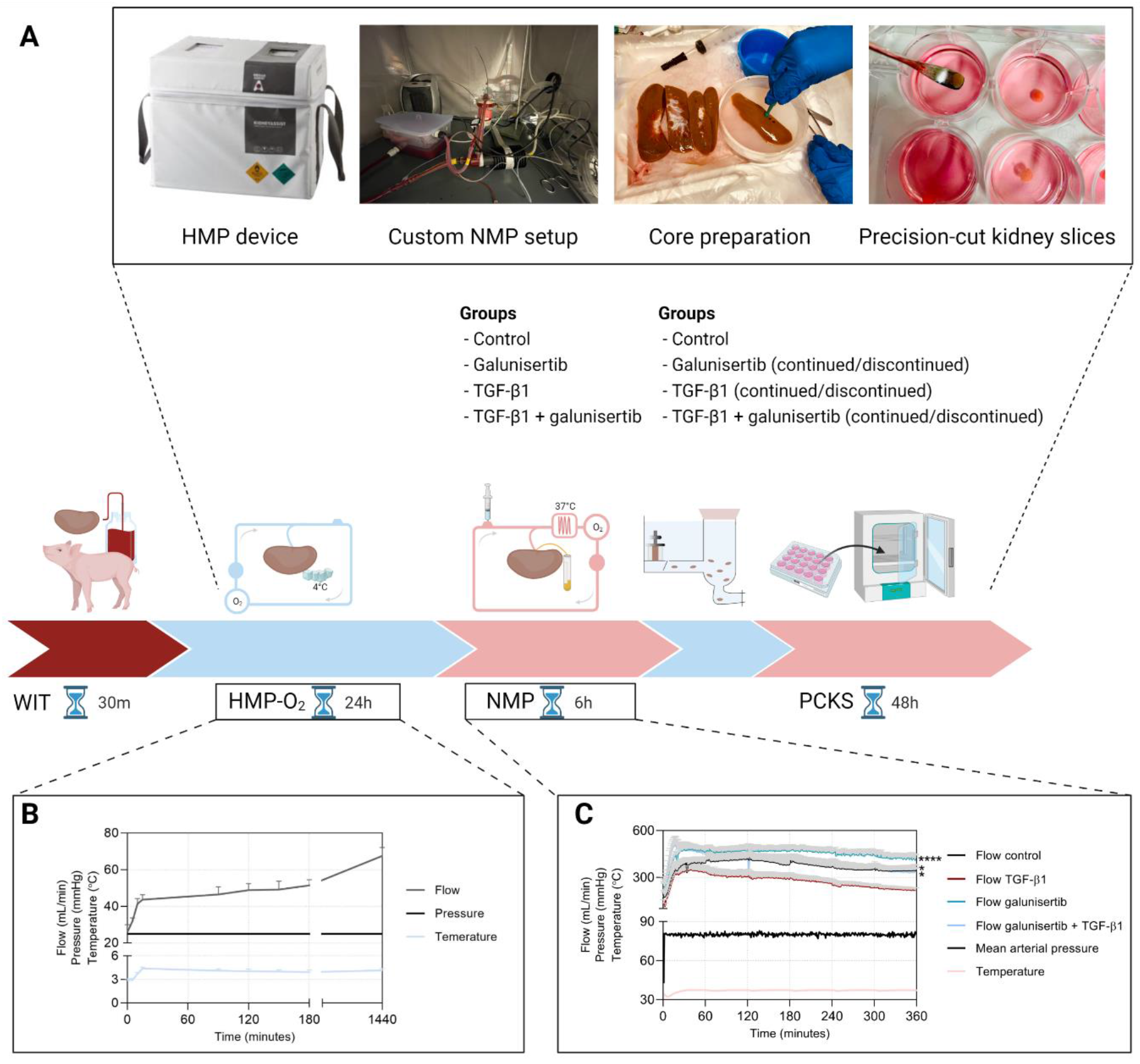
Machine perfusion and Organ slices as a Platform for *Ex vivo* Drug delivery (MOPED) workflow (**A**) porcine kidneys were subjected to 30 min of warm ischemia, 24 h of oxygenated HMP, and 6 h of NMP, after which slices were prepared from the respective kidneys, (**B**) perfusion parameters during oxygenated HMP (*n* = 32), and (**C**) perfusion parameters during NMP (*n* = 8 per group). * *p* < 0.05, **** *p* < 0.0001. Values are expressed as mean ± SEM. HMP = hypothermic machine perfusion, NMP = normothermic machine perfusion, WIT = warm ischemia time, PCKS = precision-cut kidney slices, TGF-β1 = transforming growth factor beta 1.

### Galunisertib promoted cell viability during NMP and in slices

Using our MOPED technique, we first examined whether TGF-β1, galunisertib, or a combination thereof compromised cell viability during NMP or in slices (Figure 2). To that end, we analyzed oxygen consumption and adenosine triphosphate (ATP) levels as well as general morphological features using a Periodic-acid Schiff (PAS) staining. Potential effects on oxidative stress were explored by measuring levels of thiobarbituric acid-reactive substances (TBARS). We observed that galunisertib significantly increased oxygen consumption during the second half of NMP (Fig. 2A). ATP levels, however, were not affected during NMP but were significantly increased in slices treated with galunisertib (Fig. 2B). Blinded histological scoring of PAS-stained sections was performed (Fig. 2C), and scoring revealed that kidneys had already sustained injury after 6 h of NMP, as indicated by scores reflecting tubular necrosis (Fig. 2D) and dilation (Fig. 2E) as well as glomerular dilation (Fig. 2F). Treatments produced no adverse effects during NMP. In slices, the extent of glomerular dilation and tubular necrosis increased after an incubation of 48 h. Galunisertib, however, significantly reduced tubular necrosis, whereas TGF-β1 exacerbated tubular dilation. We detected no significant differences in TBARS levels after 6 h of NMP and in slices, regardless of treatments used (Fig. 2G).

**FIG. 2.**
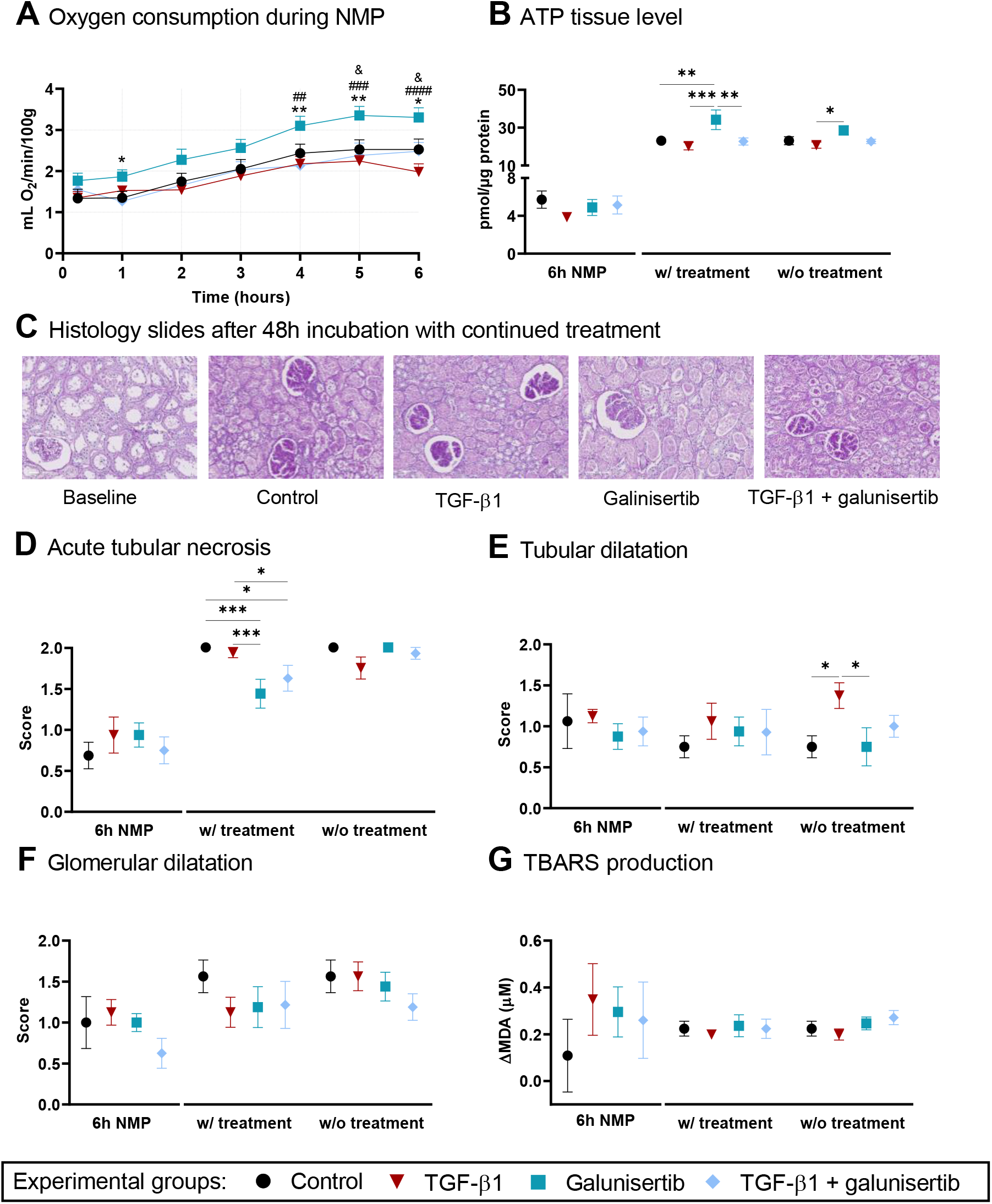
Galunisertib does not affect general viability as shown by the (**A**) oxygen consumption (#: significance between TGF-β1 and galunisertib, *: significance between galunisertib and TGF-β1 + galunisertib, and &: significance between control and galunisertib), (**B**) ATP/protein content, (**C**) general morphology as visualized using PAS staining (representative images are shown of slices after 48 h of incubation), (**D**) acute tubular necrosis scores (0 – 2), (**E**) tubular dilation scores (0 – 2), (**F**) glomerular integrity scores (0 – 2), (**G**) TBARS content in perfusate. * *p* < 0.05, ** *p* < 0.01, *** *p* < 0.001, and **** *p* < 0.0001. Values are expressed as mean ± SEM (*n* = 8). ATP = adenosine triphosphate, NMP = normothermic machine perfusion, PAS = Periodic-acid Schiff, TBARS = thiobarbituric acid reactive substances, TGF-β1 = transforming growth factor beta 1

### Galunisertib did not negatively affect renal function

We subsequently assessed whether renal function was affected by the tested treatments (Figure 3). At various timepoints during NMP, samples were collected and analyzed to determine urine production, creatinine clearance, fractional sodium excretion, and metabolic coupling. We observed that there were no significant differences in urine production between the groups, and urine production also remained stable over time, with only a minor and temporary decline at 1 h (Fig. 3A). In comparison to the control, kidneys treated with only TGF-β1 seemed to have a reduced clearance of creatinine, albeit not significantly (Fig. 3B). The fractional sodium excretion after 1 h of NMP, however, was significantly different between TGF-β1 only and TGF-β1 + galunisertib group (Fig. 3C). In addition, the fractional sodium excretion declined during the first hour of NMP, after which it remained stable. With respect to metabolic coupling, we observed no significant differences between controls and treatments (Fig. 3D).

**FIG. 3.**
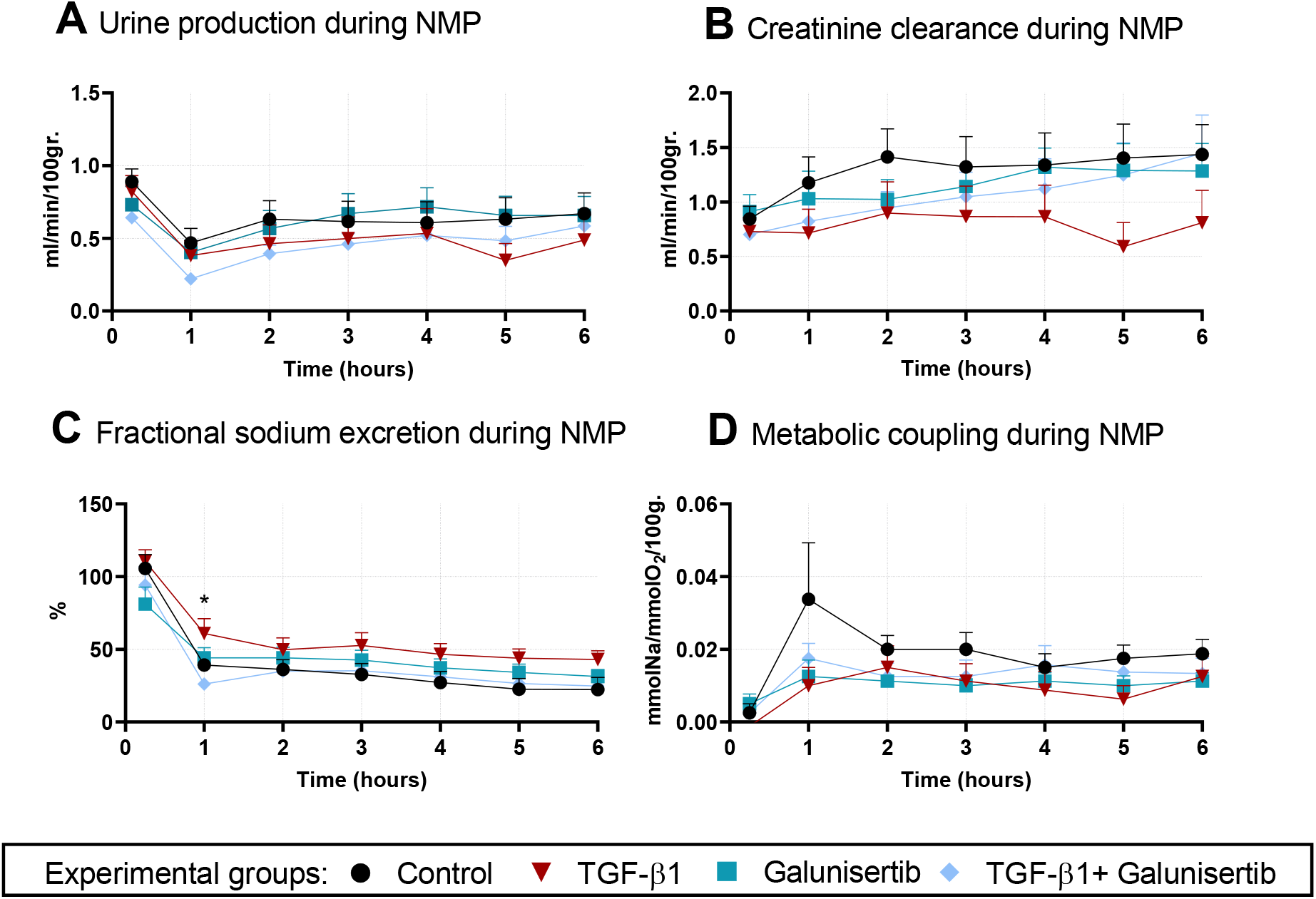
Renal function during normothermic machine perfusion as shown by (**A)** urine production rate, (**B**) creatinine clearance, (**C**) fractional sodium excretion, and (**D**) metabolic coupling. * *p* < 0.05 between TGF-β1 and TGF-β1 + galunisertib. Values are expressed as mean ± SEM (*n* = 8). NMP = normothermic machine perfusion, TGF-β1 = transforming growth factor beta 1.

### Galunisertib did not cause additional damage during NMP

To establish whether treatments caused additional damage during NMP, we measured lactate dehydrogenase (LDH) and aspartate aminotransferase (ASAT) levels in perfusate as well as total protein and *N*-acetyl-β-d-glucosaminidase (NAG) content in urine (Figure 4). The release of LDH into perfusate, which marks cell damage, increased over time, leveling off around the 3 h timepoint, but remained unaffected by TGF-β1, galunisertib, or a combination thereof (Fig. 4A). Similar observations were made for ASAT levels in the perfusate, which were used as a marker for mitochondrial damage (Fig. 4B). The total amount of excreted protein, indicating proteinuria, increased almost linearly over time and remained unaffected by the tested treatments (Fig. 4C). Likewise, the total amount of urinary NAG, indicating tubular injury, increased during NMP and was not significantly different among the experimental groups.

**FIG 4.**
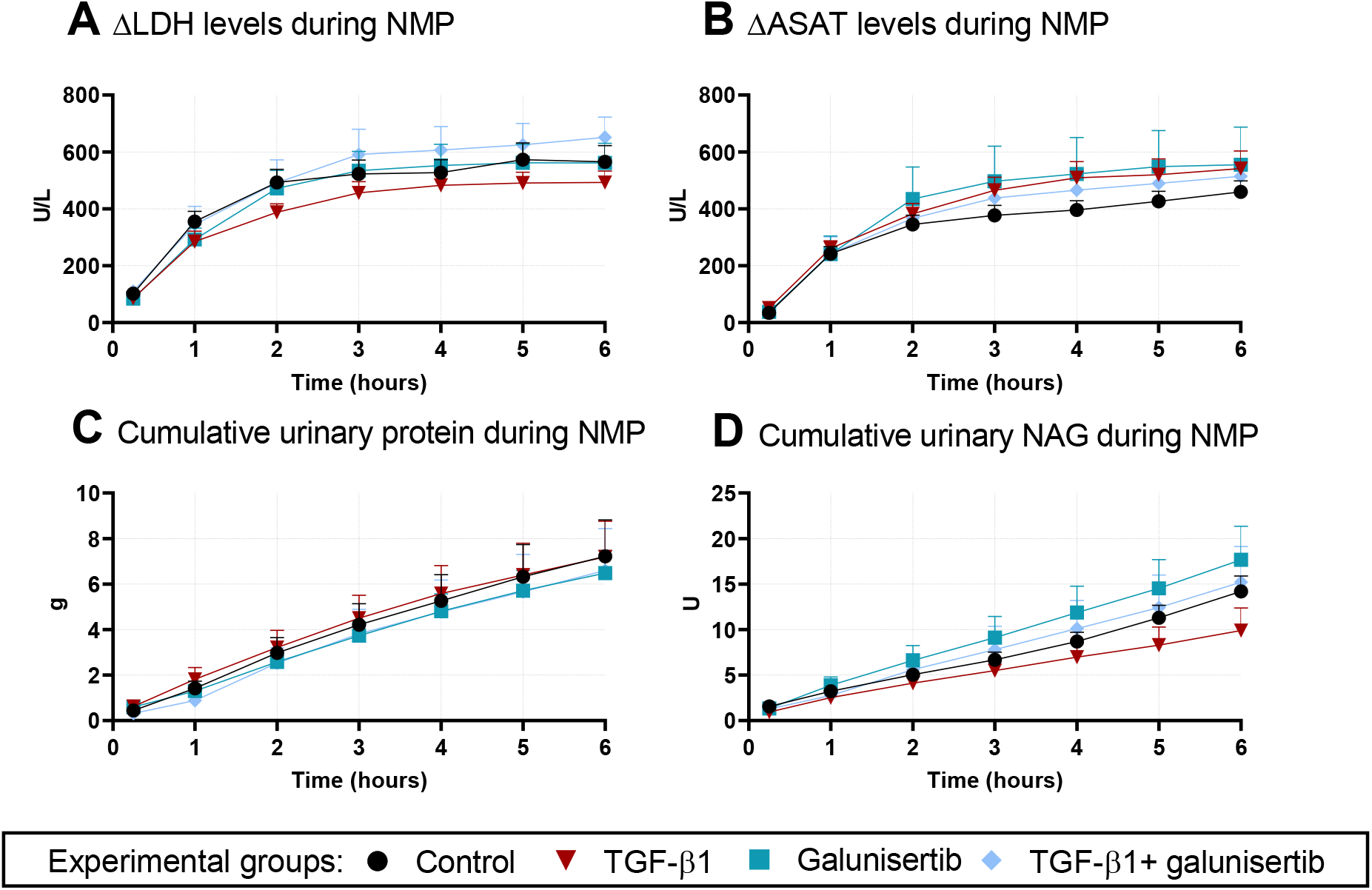
Kidney injury during NMP not aggravated by Galunisertib. As shown by (**A**) ΔLDH levels in perfusate, (**B**) ΔASAT levels in perfusate, (**C**) total urinary protein, and (**D**) total urinary NAG excreted. Values are expressed as the mean ± SEM (*n* = 8). ASAT = aspartate aminotransferase, LDH = lactate dehydrogenase, NMP = normothermic machine perfusion, TGF-β1 = transforming growth factor beta 1, uNAG = urinary *N*-acetyl-β-d-glucosaminidase.

### Galunisertib affected inflammation during NMP and in slices

After confirming that galunisertib did not affect cell viability and renal function, we analyzed mRNA expression of *IL1B*, *TGFB1*, *TNF*, and *IL6* as well as IL-6 protein content to identify potential effects on inflammation (Figure 5). We found that *IL1B* mRNA expression in kidneys treated with TGF-β1 and galunisertib was significantly lower after NMP than those treated with TGF-β1 (Fig. 5A). As for slices, we observed no treatment-dependent effects on *IL1B* mRNA expression. Similar observations were made for *TGFB1* mRNA expression, which was significantly reduced after 6 h of NMP when treated with galunisertib (Fig. 5B). This effect was maintained in slices when continuing treatments. Upon ceasing treatments, these effects diminished. Effects on *TNF* mRNA expression were also detected after NMP; galunisertib significantly increased its expression, whereas a combination of TGF-β1 and galunisertib reduced its expression (Fig. 5C). Slices showed the same pattern when treatments were continued. After 6 h of NMP, *IL6* mRNA expression in kidneys treated with TGF-β1 and galunisertib was significantly lower than those treated with TGF-β1 alone (Fig. 5D). *IL6* mRNA expression in slices remained unchanged. These effects were observed on protein level as well, as shown by the release of IL-6 into the perfusate (Fig. 5E).

**FIG 5.**
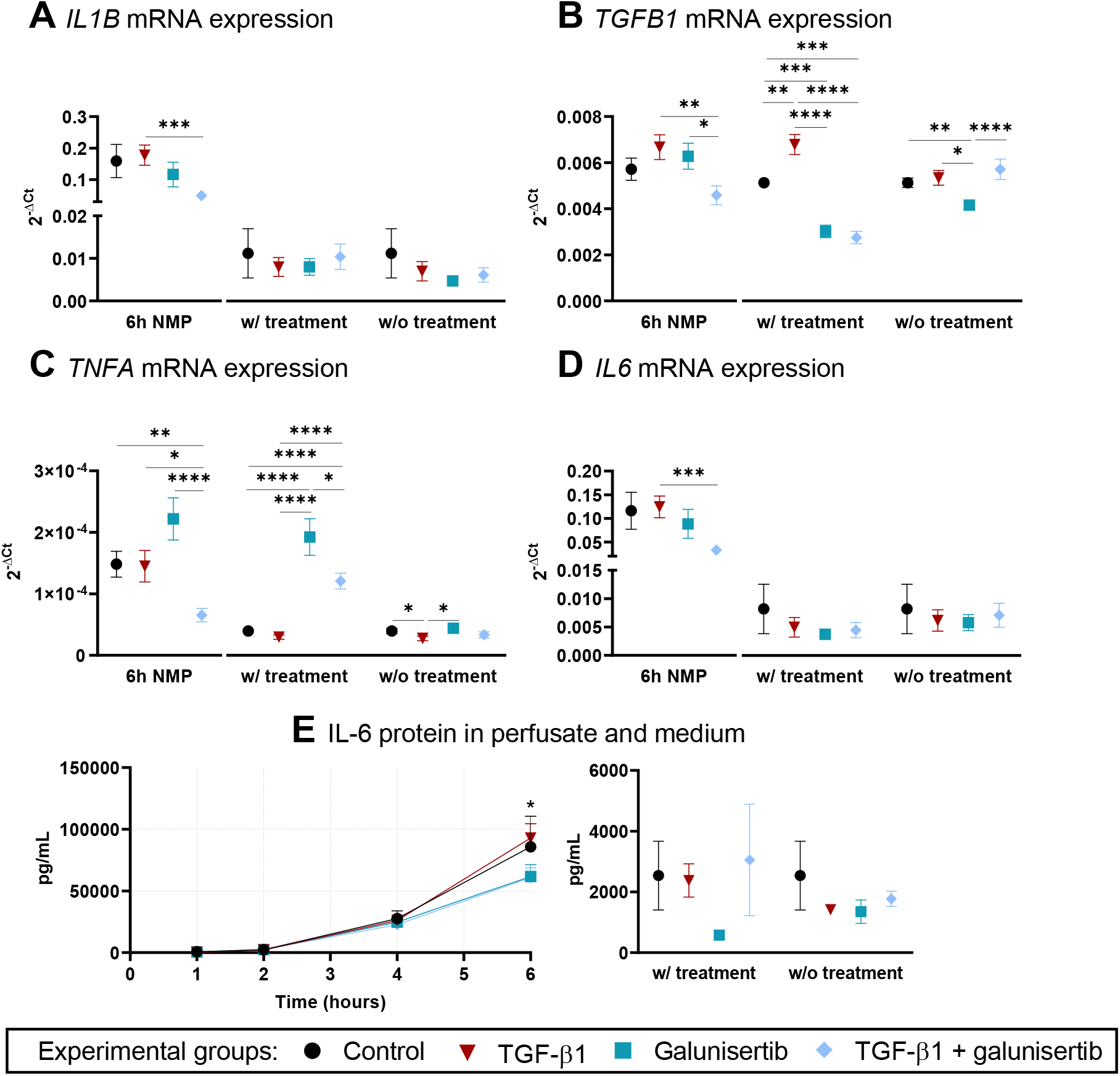
Inflammation during NMP and in slices as shown by (**A**) *IL1B* mRNA expression, (**B**) *TGFB1* mRNA expression, (**C**) *TNFA* mRNA expression, (**D**) *IL6* mRNA expression, and (**E**) IL-6 protein levels in perfusate. * *p* < 0.05 between TGF-β1 and TGF-β1 + galunisertib. Values are expressed as the mean ± SEM (*n* = 8). * *p* < 0.05, ** *p* < 0.01, *** *p* < 0.001, and **** *p* < 0.0001 IL-6 = interleukin 6, NMP = normothermic machine perfusion, TGF-β1 = transforming growth factor beta 1

### Galunisertib attenuated fibrogenesis during NMP and in slices

To determine whether galunisertib attenuated fibrogenesis—our main research question—we analyzed mRNA expression of *ACTA2*, *COL1A2*, *FN1*, *SERPINE1*, and *SERPINH1* (Figure 6). The expression of *ACTA2* mRNA was not affected by treatments after 6 h of NMP (Fig. 6A). In slices, however, mRNA expression of *ACTA2* was significantly reduced when treated with either galunisertib only or TGF-β1 and galunisertib. The opposite was detected in slices treated with TGF-β1, wherein *ACTA2* mRNA expression significantly increased. These effects were diminished when treatments were ceased in slices. Similar trends were observed for *COL1A2* (Fig. 6B) and *FN1* mRNA expression (Fig. 6C) in slices treated with galunisertib. Interestingly, *SERPINE1* mRNA expression was significantly reduced in kidneys perfused with galunisertib-containing solutions after 6 h of NMP already (Fig. 6D). Slices responded the same way when treatments were continued. With respect to *SERPINH1* mRNA expression, reductions were only observed in slices after continuing galunisertib treatments (Fig. 6E).

**FIG. 6.**
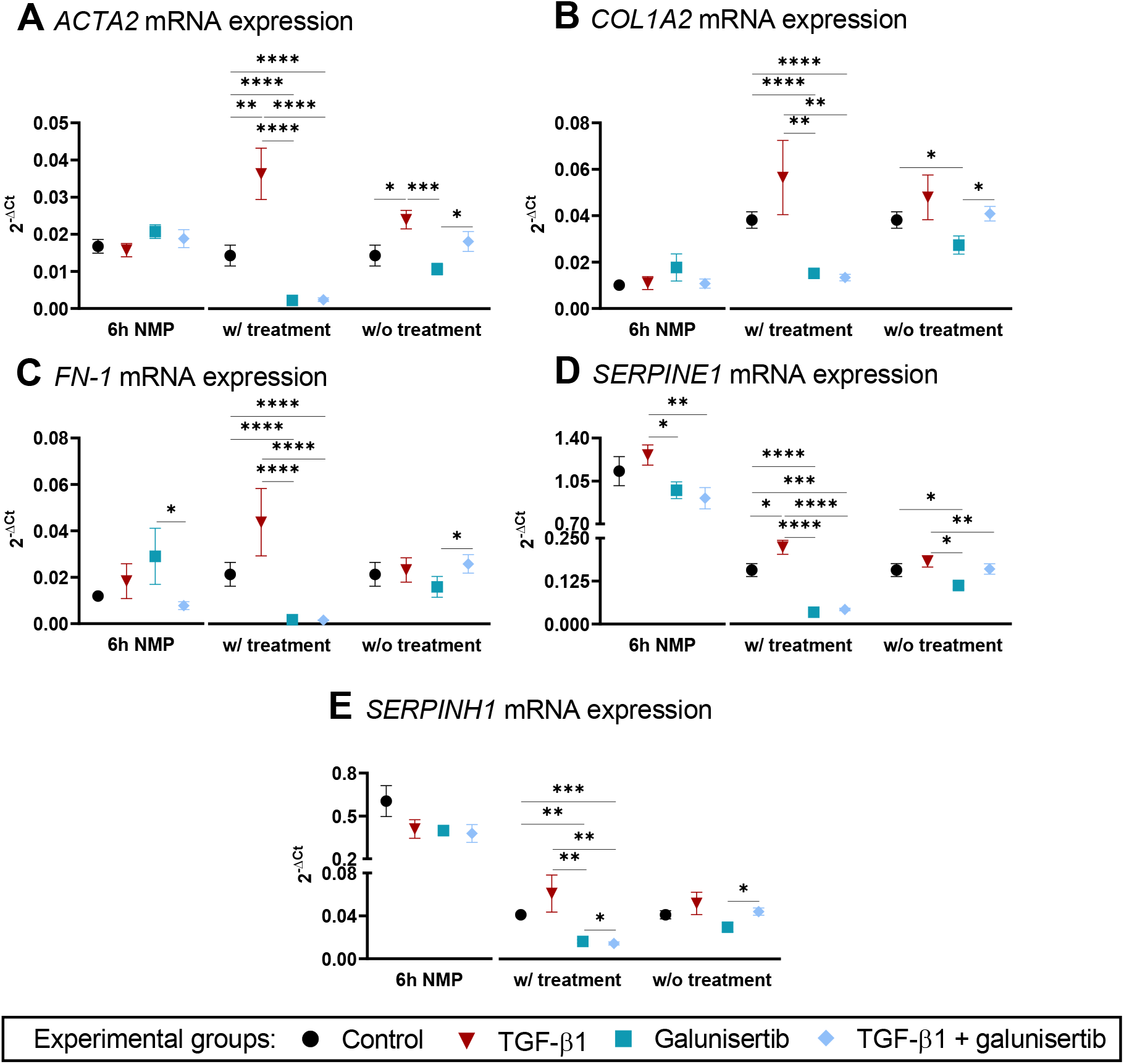
Galunisertib attenuates fibrogenesis during NMP and in slices. As shown by (**A**) *ACTA2* mRNA expression, (**B**) *COL1A2* mRNA expression, (**C**) *FN-1* mRNA expression, (**D**) *SERPINE1* mRNA expression, and (**E**) *SERPINH1* mRNA expression. * *p* < 0.05, ** *p* < 0.01, *** *p* < 0.001, and **** *p* < 0.0001. Values are expressed as the mean ± SEM (*n* = 8). NMP = normothermic machine perfusion, TGF-β1 = transforming growth factor beta 1

## DISCUSSION

Interstitial fibrosis and tubular atrophy are the main causes for long-term graft loss, and a great burden for renal transplant recipients *(10–12)*. There are currently no safe and effective therapies for halting the onset of fibrosis in kidney allografts. We therefore introduced a novel approach for studying kidney fibrosis, combining normothermic machine perfusion and precision-cut kidney slice techniques, spiked with TGF-β, called MOPED. This unique *ex vivo* model captures the full complexity of a metabolically-active isolated organ, the tissue architecture as well as cell-cell and cell-matrix interactions, which are paramount to consider in fibrosis research *(13)*. Using this newly developed model, we demonstrated that galunisertib suppresses the onset of fibrosis in kidney allografts without compromising renal viability, functionality, and injury as assessed by oxygen consumption, tissue ATP levels, histological structure, lipid peroxidation, urine production, proteinuria, creatinine clearance, fractional sodium excretion, metabolic coupling, urinary NAG, LDH, and ASAT levels.

Our main aim was to assess the early antifibrotic effects of galunisertib. Therefore, we looked at mRNA expression of key fibrosis related proteins as mRNA upregulation generally occurs sooner than protein upregulation. Galunisertib exhibited clear attenuation of *TGFB1*, *ACTA2*, *COL1A2*, *FN-1*, *SERPINE1*, and *SERPINH1,* genes encoding for TGB-β, α-SMA, FN-1, COL1A2, HSP47, and PAI-1. During fibrogenesis, TGF-β triggers a whole cascade of processes such as α-SMA expression by activated myofibroblasts with the aim of restoring tissue integrity by producing and secreting extracellular matrix (ECM) proteins, especially collagens and fibronectins *(14)*, and secretion of the collagen chaperone HSP47 *(15)*. Similar effects on gene expression of fibrosis markers were observed in previous studies using human and murine tissue slices *(16, 17)*, although the kidneys described in these studies were not subjected to NMP.

Galunisertib is a potent inhibitor of the TGF-β1 signaling pathway *(7)*. One of the main concerns of systemic inhibition of TGF-β signaling is the interference with beneficial biological processes *(18)*. We therefore tried eliminating these effect by performing *ex vivo* perfusion. As also observed in a previous study, we detected higher expression of TNF-α gene expression after treatment with galunisertib *(19)*. However, galunisertib combined with TGF-β treatment reduced TNF-α expression. We observed significantly lower IL-6 levels after 6 h NMP on both mRNA and protein level, as also observed in a previous study *(20)*, suggesting anti-inflammatory effects. Additionally, galunisertib did not compromise renal function and viability, or led to significant higher levels of urinary NAG *(21)*.

The galunisertib concentration used,10 μM, is in line with other *ex vivo* studies *(16, 17, 22)*. *In vivo*, a much lower dosage, resulting in a C_max_ of 2-6 μM, is administered to limit systemic side effects *(23, 24)*. By treating an isolated kidney, these adverse effects are circumvented and therefore a higher concentration can be used. To our knowledge, galunisertib, or any other antifibrotic molecule, has never been administered to an isolated metabolically-active kidney before.

The disadvantage of using an abattoir model is that assessment of renal function beyond NMP is not possible as the kidney cannot be (auto)transplanted. Nevertheless, experimental NMP models are commonly used platforms for optimizing the resuscitation technique itself, and *ex vivo* renal function accurately reflects the condition of the kidney *(25–27)*. In contrast to renal function, we were able to assess renal metabolism for an additional 48 hours. Kidneys are great consumers of oxygen to fuel oxidative phosphorylation and to produce ATP for the active reabsorption of sodium in the tubular cells *(28–30)*. Therefore, the renal metabolic state is a valid representation of kidney health *(31)*. Surprisingly, galunisertib resulted in a higher oxygen consumption during NMP and significantly higher ATP levels after 48 h incubation. Elevated ATP levels have not been previously observed in slices *(16, 17, 22)*. It remains unclear whether these high ATP levels are due to improved cellular respiration during NMP, or the fact that galunisertib is a competitive inhibitor for ATP-binding site of the TGF-β receptor 1 *(24)*.

We envision that our MOPED technique could be used for testing a broad range of small molecules to ultimately improve transplant outcomes. Pretreatment with TGF-β is an effective method to induce the onset of fibrosis in a relative short period of time. Additionally, the great advantage of using porcine kidneys obtained from the abattoir is that they are, genetically, physiologically and heterogeneity wise, very similar to human kidneys *(32)*, and limit the use of laboratory animals. As a follow-up to the porcine models, donated human kidneys rejected for transplantation could be used, leading to more translatable results.

In summary, this study not only introduces MOPED, but also a novel therapeutic approach to target one of the main burdens in renal transplantation. With further research using transplant models, discarded human kidneys and clinical evaluation, we envision that small molecule drugs such as galunisertib could provide the aid to renal allograft related interstitial fibrosis and tubular atrophy.

## METHODS

### Study design

This study aimed to investigate the antifibrotic effect of galunisertib using a combination of normothermic machine perfusion and precision-cut kidney slices. Extensive pilot studies were performed to optimize our experimental model, study design, and protocol. Exclusion criteria were: visibly damaged kidneys (cuts, cysts, etc.) or kidneys with aberrant arteries during organ retrieval, insufficient kidney function and technical issues during NMP, and infections during slice incubation. These criteria were established prospectively. Heterogeneity amongst animals was expected as the pigs were not bred in a standardized manner. Therefore, the sample size of n=8 per experimental group was determined based on a power calculation and was not altered during the study. Only one kidney from each pig was used and randomized into experimental groups. Unpaired statistical analyses were therefore applied. Due to the heterogeneity, outliers were not excluded. Our primary endpoint was the effect of galunisertib on gene expression of fibrosis related markers. Investigators were not blinded during the execution of NMP and slice experiments, however they were blinded during the execution of all consecutive analyses.

### Organ procurement and preservation

Kidneys were obtained from female Dutch Landrace pigs from a local abattoir in accordance with all guidelines of the Dutch food safety authority. Pigs were anesthetized by an electrical shock and instantly exsanguinated. After 30 min of warm ischemia, the kidneys were flushed with ice-cold saline. After cannulation, kidneys were connected to a hypothermic machine perfusion device (Kidney Assist Transport, XVIVO, Göteborg, Sweden). HMP was performed for 24 h, at 4 °C, using pulsatile pressure-controlled perfusion with a mean arterial pressure of 25 mmHg and 100% oxygenated (100 mL/min) University of Wisconsin machine perfusion solution (UW-MP) (Belzer MPS, Bridge to Life, London, UK).

### Normothermic machine perfusion

After HMP, kidneys were flushed with ice-cold saline and cannulated for NMP. NMP was carried out using a custom built perfusion circuit containing an organ chamber with tubing and a centrifugal pump (Medos Medizintechnik AG, Radeberg, Germany) controlled by a custom designed pressure-controlled perfusion machine (LabView Software) set at 80 mmHg with an amplitude of 15, a clinical-grade oxygenator/heat exchanger (Hilite 800 LT, Medos Medizintechnik AG) supplied with carbogen (95% O_2_/5% CO_2_) at a rate of 0.5 ml/min, a clinical-grade pressure sensor (Edwards Lifesciences) and a ultra-sensitive flow sensor (Transonic). The total setup was surrounded by a heating cabinet to keep the temperature stable at 37° C with the help of temperature sensors and an electric heater (Tristar).

The perfusate contained 835 mL of heparinized leukocyte-depleted autologous whole blood with a hematocrit of 36% and 165 mL of Ringer’s solution (Baxter, Utrecht, The Netherlands), as well as 10 μg/mL ciprofloxacin (Merck, Amsterdam, The Netherlands), 0.1% sodium bicarbonate (B. Braun, Melsungen, Germany), 0.0625% glucose (Baxter), 8.3 μg/mL dexamethasone (Centrafarm, Etten-Leur, The Netherlands), 10 μg/mL mannitol (Baxter), 0.135 μg/mL creatinine (Merck), and 2.7 μg/mL sodium nitroprusside (Merck). During NMP, the perfusate was also supplemented through continuous infusion (20 mL/h) of a solution comprising 10% aminoplasmal (B. Braun), 0.25% sodium bicarbonate (B. Braun), 0.2 U/mL insulin (Novo Nordisk, Bagsvaerd, Denmark), and 35 IE/mL heparin (Leo Pharma, Ballerup, Denmark). After 1 h of NMP, treatments were started by exposing kidneys to 5 ng/mL TGF-β (Sigma Aldrich, Amsterdam, The Netherlands), 10 μM galunisertib (Axon Medchem, Groningen, The Netherlands), or a combination thereof. The perfusate of control kidneys was supplemented with DMSO, which served as a vehicle. Urine, perfusate, and biopsies were sampled at various timepoints during NMP. Urine was recirculated after sampling.

### Precision-cut kidney slices

After NMP, the kidneys were immediately flushed with ice-cold saline. Cortical tissue cores were subsequently prepared using a biopsy puncher. The tissue cores were transferred to ice-cold UW cold storage solution (Bridge-to-Life). Slices with a thickness of 300 μm and a diameter of 6 mm were prepared with a Krumdieck tissue slicer (Alabama Research and Development, Munford, USA), as described previously *(17)*. Slices were cultured in 12-well plates, containing pre-warmed (37 °C) culture medium (1.3 mL/well), at 5% CO_2_ and 80% O_2_ while being gently shaken (90 cycles/min). Culture medium comprised William’s Medium E + GlutaMAX (Invitrogen, Landsmeer, The Netherlands), 10 μg/mL ciprofloxacin (Invitrogen), and 0.25 μg/mL amphotericin B (Invitrogen). To determine whether effects persisted upon ceasing or continuing treatments, slices were cultured for 48 h with 5 ng/mL TGF-β1 (Sigma Aldrich), 10 μM galunisertib (Axon Medchem), or a combination thereof; slices cultured in DMSO (vehicle) served as a control group. Culture media, including respective treatments, were refreshed after 24 h.

### Cell viability assay

Using a Minibead-beater (2 cycles of 45 s), cortical biopsies and slices were homogenized in ice-cold sonication solution (70% ethanol and 2 mM EDTA). After centrifugation (16,000 x g at 4 °C for 5 min), supernatants were analyzed using an ATP Bioluminescence Kit (Roche Diagnostics, Mannheim, Germany). Supernatants were subsequently stored overnight at 37 °C, allowing for the evaporation of sonication solution. The respective pellets were reconstituted, and the resulting supernatants were analyzed using a Pierce BCA Protein Assay Kit (Invitrogen). ATP values were normalized to protein content.

### Lipid peroxidation assay

Culture medium and perfusate samples were analyzed to investigate the formation of TBARS, which are often used as an indicator of oxidative stress. The protocol for this analysis has been described in detail previously *(33)*.

### Evaluation of perfusion parameters

The renal flow rate and urine production were logged during NMP. Creatinine and sodium concentrations in urine and perfusate samples, LDH and ASAT in perfusate samples, and protein levels in urine samples were analyzed in a standardized manner by the clinical chemistry department of the University Medical Center Groningen. Additionally, lactate, potassium and hemoglobin content as well as partial oxygen pressure and hemoglobin saturation were measured using an ABL90 FLEX blood gas analyzer (Radiometer, Zoetermeer, The Netherlands). uNAG levels were determined as described previously *(34, 35)*. Equations for calculating oxygen consumption, metabolic coupling, creatinine clearance, and fractional sodium excretion are shown in appendix 1.

### Histological assessment

Cortical biopsies and slices were fixed in 4% formalin, after which they were dehydrated by immersing tissues in a series of ethanol solutions of increasing concentrations. The tissues were then cleared in xylene, embedded in paraffin wax, and cut into sections of 4 μm. Sections were stained using a conventional PAS staining to visualize morphological features. Sections were scanned with a C9600 NanoZoomer (Hamamatsu Photonics, Hamamatsu, Japan) to obtain high-resolution digital data. Semi-quantitative scores were assigned to PAS-stained sections in a blinded manner by three individuals, marking glomerular dilatation and structure, tubular dilatation and acute tubular necrosis (Appendix 2).

### Gene expression analysis

Total RNA was extracted from cortical biopsies and slices using TRIzol reagent (Invitrogen, Landsmeer, The Netherlands). The yield of extracted RNA was analyzed with a NanoDrop 1000 spectrophotometer (NanoDrop Technologies, Wilmington, USA), and the quality was assessed using RNA electrophoresis. Extracted RNA was reverse transcribed using M-MLV Reverse Transcriptase (Invitrogen) at 37 °C for 50 min. Real-time quantitative polymerase chain reaction (qPCR) was conducted using specific primers (Appendix 3), *Taq* DNA Polymerase (Invitrogen), and a QuantStudio 7 Flex qPCR machine (Applied Biosystems, Bleiswijk, The Netherlands), which was configured with 1 cycle of 10 min at 95 °C and 40 consecutive cycles of 15 s at 95 °C and 1 min at 60 °C. Expression levels were calculated as 2^−ΔCt^, using *ACTB* as a reference gene.

### Interleukin-6 (IL-6) immunoassay

Culture medium and perfusate samples were analyzed with a Porcine IL-6 DuoSet enzyme-linked immunosorbent assay (ELISA) (Bio-Techne, Abingdon, UK), according to the manufacturer’s instructions.

### Statistical analysis

GraphPad Prism (version 8.4.2.) was used to visualize and analyze the data. All data are expressed as mean with standard error of the mean (SEM). For longitudinal data, differences across all experimental groups were assessed using a two-way analysis of variance (ANOVA) with Geisser-Greenhouse correction followed by Tukey’s multiple comparisons. Single time points were analyzed using an one-way ANOVA followed by Fisher’s least significant difference test. All statistical tests were two-tailed, and differences between groups were considered to be statistically significant when *p* < 0.05.

## APPENDIX 1

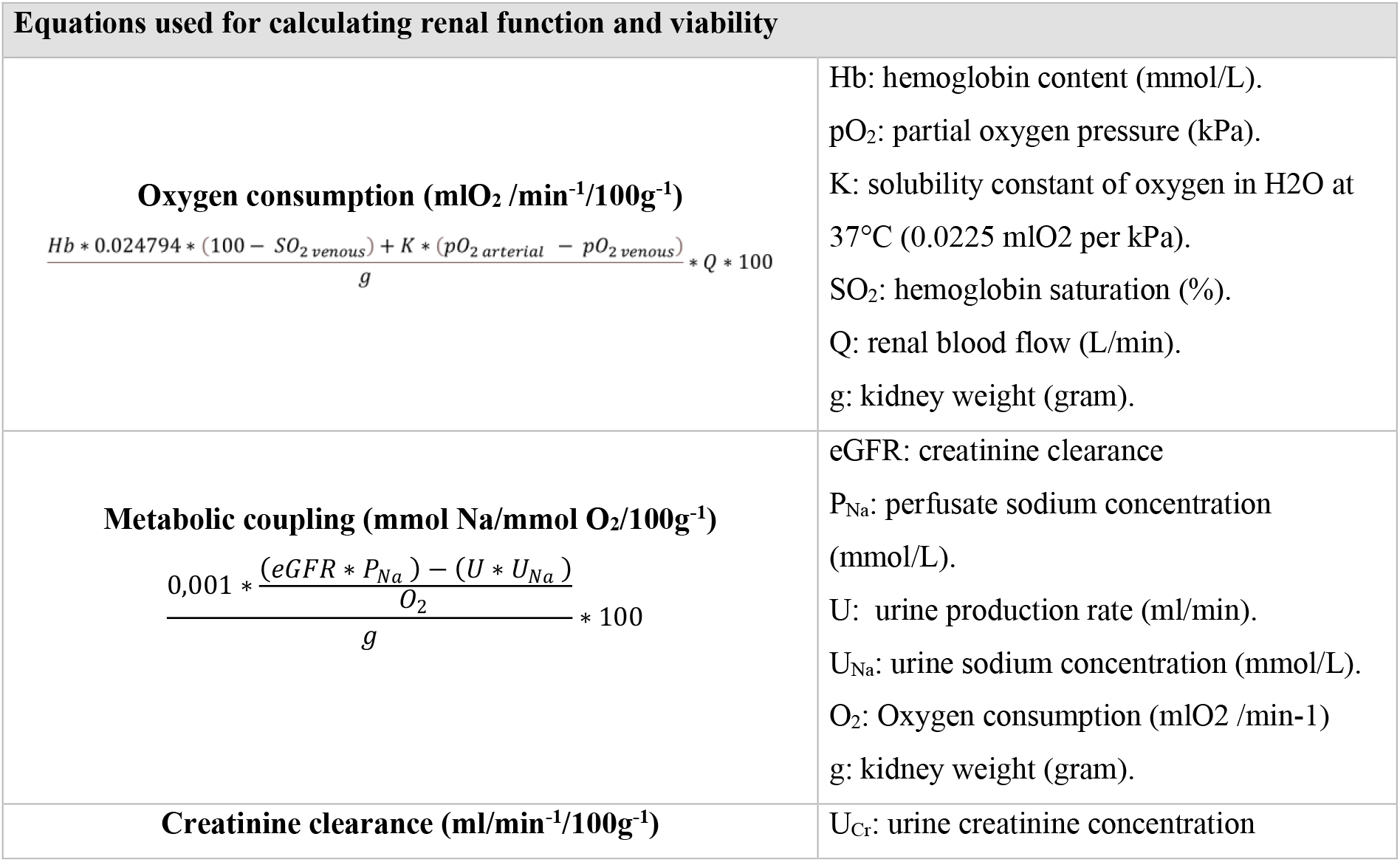
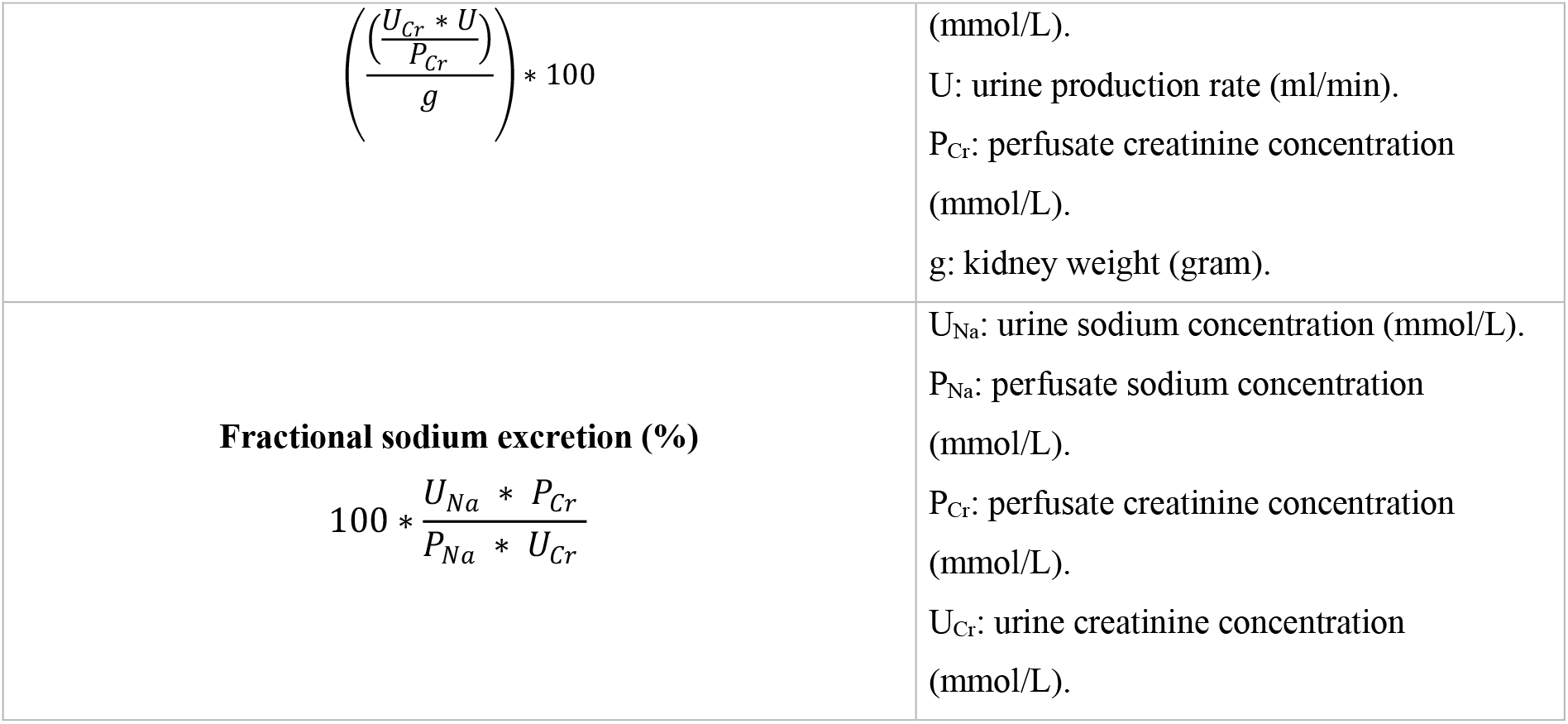

| Equations used for calculating renal function and viability |  |
| --- | --- |
| **Oxygen consumption (mlO<sub>2</sub> /min<sup>-1</sup>/100g<sup>-1</sup>)**<br>$\frac{Hb * 0.024794 * (100 - SO_{2venous}) + K * (pO_{2arterial} - pO_{2venous})}{g} * Q * 100$ | Hb: hemoglobin content (mmol/L).<br>pO<sub>2</sub>: partial oxygen pressure (kPa).<br>K: solubility constant of oxygen in H<sub>2</sub>O at 37°C (0.0225 mlO<sub>2</sub> per kPa).<br>SO<sub>2</sub>: hemoglobin saturation (%).<br>Q: renal blood flow (L/min).<br>g: kidney weight (gram). |
| **Metabolic coupling (mmol Na/mmol O<sub>2</sub>/100g<sup>-1</sup>)**<br>$\frac{0,001 * \frac{(eGFR * P_{Na}) - (U * U_{Na})}{O_2}}{g} * 100$ | eGFR: creatinine clearance<br>P<sub>Na</sub>: perfusate sodium concentration (mmol/L).<br>U: urine production rate (ml/min).<br>U<sub>Na</sub>: urine sodium concentration (mmol/L).<br>O<sub>2</sub>: Oxygen consumption (mlO<sub>2</sub> /min-1)<br>g: kidney weight (gram). |
| **Creatinine clearance (ml/min<sup>-1</sup>/100g<sup>-1</sup>)** | U<sub>Cr</sub>: urine creatinine concentration |
| $\left( \frac{\left( \frac{U_{Cr} * U}{P_{Cr}} \right)}{g} \right) * 100$ | (mmol/L).<br>U: urine production rate (ml/min).<br>P<sub>Cr</sub>: perfusate creatinine concentration (mmol/L).<br>g: kidney weight (gram). |
| **Fractional sodium excretion (%)**<br>$100 * \frac{U_{Na} * P_{Cr}}{P_{Na} * U_{Cr}}$ | U<sub>Na</sub>: urine sodium concentration (mmol/L).<br>P<sub>Na</sub>: perfusate sodium concentration (mmol/L).<br>P<sub>Cr</sub>: perfusate creatinine concentration (mmol/L).<br>U<sub>Cr</sub>: urine creatinine concentration (mmol/L). |

## APPENDIX 2

| Glomerular dilatation and structural changes |  |
| --- | --- |
| 0 | Glomeruli are still intact and demonstrate a clear clove-structure. Capillaries have thin walls and are clearly visible. |
| 1 | The clove-structure is less present, and glomeruli are not entirely intact. Capillaries are less present and dilated, and capillary walls are not clearly visible. Heightened dilatation of the Bowmans space. Bowmans's space can be filled with necrotic tissue/cells. |
| 2 | Complete dilatation of the Bowmans space (50-100% of the glomerulus' size). Glomeruli have decreased in size and are crumpled up. |
| Tubular dilatation |  |
| 0 | The tubules are not dilated. The brush border and tubular membranes are clearly visible |
| 1 | The tubules are twice their regular size. Intratubular space is visible and tubule membranes are no longer attached to each other. Tubular cells still have a regular size. The brush border is no longer visible. |
| 2 | The tubules have increased to more than twice their regular size. The tubular cells are not visible and have irregular shapes. |
| Acute tubular necrosis |  |
| 0 | No necrotic tissue visible. The tubules are not dilated. The brush border and tubular membranes are clearly visible |
| 1 | Some necrotic tissue is visible. Little thickening of the membranes, and the tubules contain some cells. |
| 2 | Vacuolization of cells, and loss of brush border. Dilation of tubular lumen, and mild interstitial inflammation. The epithelium of the proximal tubules is flattened and simplified. No clear tubular definition. Thickening of the tubule membrane and the tubules contain many cells. |

## APPENDIX 3

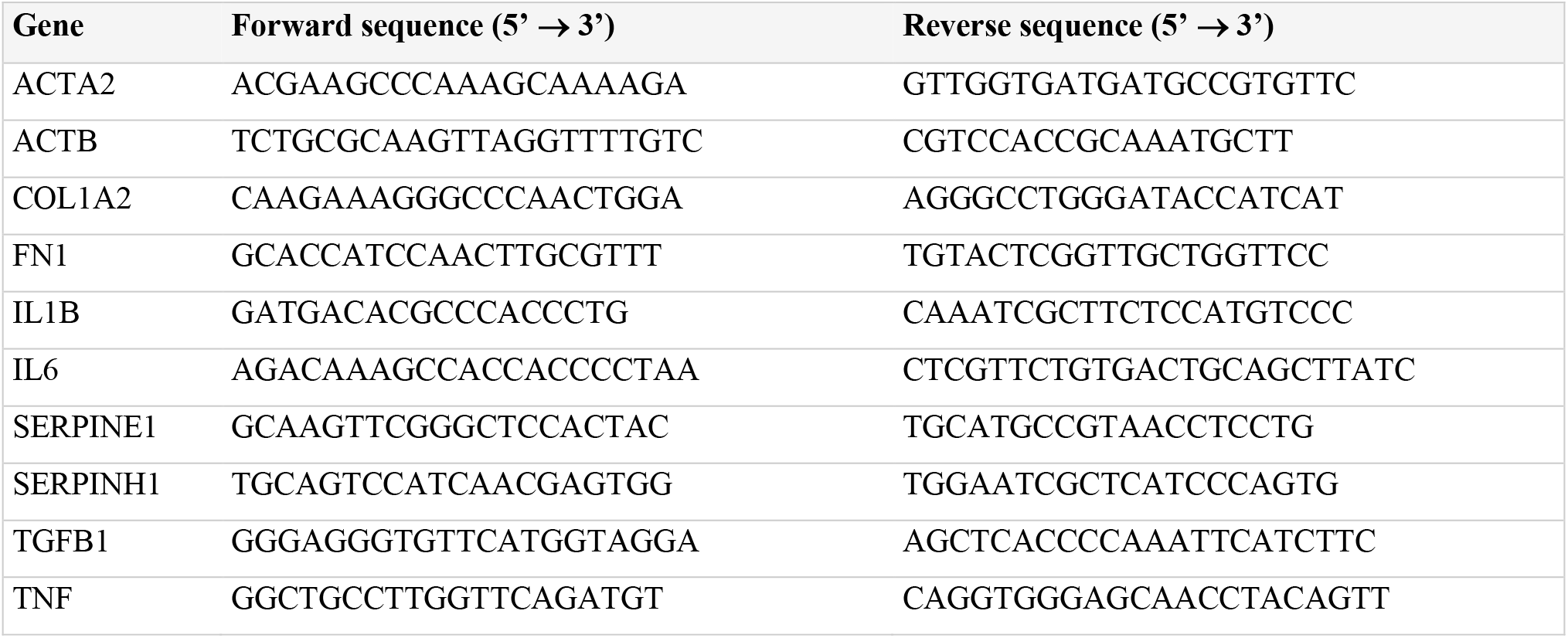

## ABBREVIATIONS

ANOVA: Analysis of variance
ASAT: Aspartate aminotransferase
ATP: Adenosine triphosphate
COL1A2: Collagen alpha-2(I) chain
ECM: Extracellular matrix
ELISA: Enzyme-linked immunosorbent assay
FN-1: Fibronectin-1
HMP: Hypothermic machine perfusion
HSP47: Heat shock protein 47
IF/TA: Interstitial fibrosis and tubular atrophy
IL1B: Interleukin-1 beta
IL-6: Interleukin-6
IRI: Ischemia-reperfusion injury
LDH: Lactate dehydrogenase
MOPED: Machine perfusion and Organ slices as a Platform for Ex vivo Drug delivery
NAG: N-acetyl-β-d-glucosaminidase
NMP: Normothermic machine perfusion
PAI-1: Plasminogen activator inhibitor 1
PAS: Periodic-acid Schiff
SEM: Standard error of the mean
TBARS: Thiobarbituric acid-reactive substances
TGF-β: Transforming growth factor beta
TNF: Tumor necrosis factor
UW-MP: University of Wisconsin machine perfusion solution
α-SMA: Actin, aortic smooth muscle

## ACKNOWLEDGEMENTS

**General**: The authors are very grateful for abattoir Kroon Vlees for their collaboration and providing the kidneys for this research. Many thanks to Yvette Jansen, Danique Zantinge, Meindert Tangerman, Petra Ottens and Janneke Wiersema-Buist for their assistance with execution of the experiments and analyses.

## Author contributions

Conceptualization: LLL, HGDL, PO, MJRR; Execution of experiments and analyses: LLL; Data analysis and visualization: LLL; Supervision: HGDL, BMK, PO; Writing – original draft: LLL, MJRR; Writing – review & editing: LLL, HGDL, BMK, PO, MJRR.

## Data and materials availability

All additional data is available upon request.

## REFERENCES

1. T. Vanhove, R. Goldschmeding, D. Kuypers, Kidney Fibrosis, Transplantation 101, 713–726 (2017).

2. L. L. van Leeuwen, H. G. D. Leuvenink, P. Olinga, M. J. R. Ruigrok, Shifting Paradigms for Suppressing Fibrosis in Kidney Transplants: Supplementing Perfusion Solutions With Anti-fibrotic Drugs, Front. Med. 0, 2917 (2022).

3. T. Saritas, R. Kramann, Kidney Allograft Fibrosis: Diagnostic and Therapeutic Strategies, Transplantation 105, E114–E130 (2021).

4. C. Moers, J. M. Smits, M.-H. J. Maathuis, J. Treckmann, F. Van Gelder, B. P. Napieralski, M. Van Kasterop-Kutz, J. J. Homan Van Der Heide, J.-P. Squifflet, E. Van Heurn, G. R. Kirste, A. Rahmel, H. G. D. Leuvenink, A. Paul, J. Pirenne, R. J. Ploeg, Machine Perfusion or Cold Storage in Decreased-Donor Kidney Transplantation, New Engl. J. Med. 360, 7–19 (2009).

5. I. Jochmans, C. Moers, J. M. Smits, H. G. D. Leuvenink, J. Treckmann, A. Paul, A. Rahmel, J.-P. Squifflet, E. van Heurn, D. Monbaliu, R. J. Ploeg, J. Pirenne, Machine Perfusion Versus Cold Storage for the Preservation of Kidneys Donated After Cardiac Death, Ann. Surg. 252, 756–764 (2010).

6. S. A. Hosgood, E. Thompson, T. Moore, C. H. Wilson, M. L. Nicholson, Normothermic machine perfusion for the assessment and transplantation of declined human kidneys from donation after circulatory death donors, Br. J. Surg. 105, 388–394 (2018).

7. Y. Isaka, Targeting TGF-β Signaling in Kidney Fibrosis., Int. J. Mol. Sci. 19 (2018).

8. E. Bigaeva, E. G. D. Stribos, H. A. M. Mutsaers, B. Piersma, A. M. Leliveld, I. J. de Jong, R. A. Bank, M. A. Seelen, H. van Goor, L. Wollin, P. Olinga, M. Boersema, Inhibition of tyrosine kinase receptor signaling attenuates fibrogenesis in an ex vivo model of human renal fibrosis, Am. J. Physiol. Physiol. 318, F117–F134 (2020).

9. M. S. Jensen, H. A. M. Mutsaers, S. J. Tingskov, M. Christensen, M. G. Madsen, P. Olinga, T. H. Kwon, R. Nørregaard, Activation of the prostaglandin E2 EP2 receptor attenuates renal fibrosis in unilateral ureteral obstructed mice and human kidney slices, Acta Physiol. 227, 13291 (2019).

10. F. G. Cosio, J. P. Grande, T. S. Larson, J. M. Gloor, J. A. Velosa, S. C. Textor, M. D. Griffin, M. D. Stegall, Kidney Allograft Fibrosis and Atrophy Early After Living Donor Transplantation, Am. J. Transplant. 5, 1130–1136 (2005).

11. P. Dinis, P. Nunes, L. Marconi, F. Furriel, B. Parada, P. Moreira, A. Figueiredo, C. Bastos, A. Roseiro, V. Dias, F. Rolo, F. Macário, A. Mota, Kidney Retransplantation: Removal or Persistence of the Previous Failed Allograft?, Transplant. Proc. 46, 1730–1734 (2014).

12. A. J. Matas, K. J. Gillingham, A. Humar, R. Kandaswamy, D. E. R. Sutherland, W. D. Payne, T. B. Dunn, J. S. Najarian, 2202 kidney transplant recipients with 10 years of graft function: what happens next?, Am. J. Transplant 8, 2410–9 (2008).

13. N. G. Frangogiannis, Fibroblast—Extracellular Matrix Interactions in Tissue Fibrosis, Curr. Pathobiol. Rep. 4, 11 (2016).

14. V. LeBleu, G. Taduri, J. O’Connell, Y. Teng, V. Cooke, C. Woda, H. Sugimoto, R. Kalluri, Origin and function of myofibroblasts in kidney fibrosis, Nat. Med. 19, 1047–1053 (2013).

15. M. S. Razzaque, V. T. Le, T. Taguchi, Heat Shock Protein 47 and Renal Fibrogenesis, Cell. Stress Responses Ren. Dis. 148, 57–69 (2005).

16. E. Bigaeva, E. Gore, H. A. M. Mutsaers, D. Oosterhuis, Y. O. Kim, D. Schuppan, R. A. Bank, M. Boersema, P. Olinga, Exploring organ-specific features of fibrogenesis using murine precision-cut tissue slices, Biochim. Biophys. Acta - Mol. Basis Dis. 1866 (2020).

17. E. Bigaeva, N. P. Cavanzo, E. G. D. Stribos, A. J. de Jong, C. Biel, H. A. M. Mutsaers, M. S. Jensen, R. Nørregaard, A. M. Leliveld, I. J. de Jong, J. L. Hillebrands, H. van Goor, M. Boersema, R. A. Bank, P. Olinga, Predictive value of precision-cut kidney slices as an ex vivo screening platform for therapeutics in human renal fibrosis, Pharmaceutics 12, 459 (2020).

18. C. Y. Huang, C. L. Chung, T. H. Hu, J. J. Chen, P. F. Liu, C. L. Chen, Recent progress in TGF-β inhibitors for cancer therapy, Biomed. Pharmacother. 134, 111046 (2021).

19. S. K. Hira, A. Rej, A. Paladhi, R. Singh, J. Saha, I. Mondal, S. Bhattacharyya, P. P. Manna, Galunisertib Drives Treg Fragility and Promotes Dendritic Cell-Mediated Immunity against Experimental Lymphoma, iScience 23, 101623 (2020).

20. X. Liu, M. Yu, Y. Chen, J. Zhang, Galunisertib (LY2157299), a transforming growth factor-β receptor I kinase inhibitor, attenuates acute pancreatitis in rats, Brazilian J. Med. Biol. Res. = Rev. Bras. Pesqui. medicas e Biol. 49 (2016).

21. R. G. Price, The role of NAG (N-acetyl-beta-D-glucosaminidase) in the diagnosis of kidney disease including the monitoring of nephrotoxicity - PubMed, Clin. Nephrol. 38, 9–14 (1992).

22. T. Luangmonkong, S. Suriguga, E. Bigaeva, M. Boersema, D. Oosterhuis, K. P. de Jong, D. Schuppan, H. A. M. Mutsaers, P. Olinga, Evaluating the antifibrotic potency of galunisertib in a human *ex vivo* model of liver fibrosis, Br. J. Pharmacol. 174, 3107–3117 (2017).

23. J. Rodón, M. Carducci, J. M. Sepulveda-Sánchez, A. Azaro, E. Calvo, J. Seoane, I. Braña, E. Sicart, I. Gueorguieva, A. Cleverly, N. S. Pillay, D. Desaiah, S. T. Estrem, L. Paz-Ares, M. Holdhoff, J. Blakeley, M. M. Lahn, J. Baselga, Pharmacokinetic, pharmacodynamic and biomarker evaluation of transforming growth factor-β receptor I kinase inhibitor, galunisertib, in phase 1 study in patients with advanced cancer, Invest. New Drugs 33, 357–370 (2015).

24. J. M. Yingling, W. T. McMillen, L. Yan, H. Huang, J. S. Sawyer, J. Graff, D. K. Clawson, K. S. Britt, B. D. Anderson, D. W. Beight, D. Desaiah, M. M. Lahn, K. A. Benhadji, M. J. Lallena, R. B. Holmgaard, X. Xu, F. Zhang, J. R. Manro, P. W. Iversen, C. V. Iyer, R. A. Brekken, M. D. Kalos, K. E. Driscoll, Preclinical assessment of galunisertib (LY2157299 monohydrate), a first-in-class transforming growth factor-β receptor type I inhibitor, Oncotarget 9, 6659 (2018).

25. L. H. Venema, A. Brat, C. Moers, N. A. ’t Hart, R. J. Ploeg, P. Hannaert, T. Minor, A. H. G. D. Leuvenink, Effects of Oxygen During Long-term Hypothermic Machine Perfusion in a Porcine Model of Kidney Donation After Circulatory Death, Transplantation 103, 2057–2064 (2019).

26. L. H. Venema, L. L. van Leeuwen, R. A. Posma, H. van Goor, R. J. Ploeg, P. Hannaert, T. Hauet, T. Minor, H. G. D. Leuvenink, Impact of Red Blood Cells on Function and Metabolism of Porcine Deceased Donor Kidneys During Normothermic Machine Perfusion, Transplantation (2021).

27. A. Weissenbacher, L. Lo Faro, O. Boubriak, M. F. Soares, I. S. Roberts, J. P. Hunter, D. Voyce, N. Mikov, A. Cook, R. J. Ploeg, C. C. Coussios, P. J. Friend, Twenty-four-hour normothermic perfusion of discarded human kidneys with urine recirculation, Am. J. Transplant 19, 178–192 (2019).

28. R. S. Balaban, Regulation of oxidative phosphorylation in the mammalian cell, Am. J. Physiol. - Cell Physiol. 258 (1990).

29. N. A. Lassen, U. Lassen, O. Munck, J. H. Thaysen, Oxygen consumption and sodium reabsorption by the kidney. Discussion of a theory., Presse Med. 69, 1259–1260 (1961).

30. P. Singh, S. E. Ricksten, G. Bragadottir, B. Redfors, L. Nordquist, Renal oxygenation and haemodynamics in acute kidney injury and chronic kidney disease, Clin. Exp. Pharmacol. Physiol. 40, 138–47 (2013).

31. M. I. Bellini, J. Yiu, M. Nozdrin, V. Papalois, The Effect of Preservation Temperature on Liver, Kidney, and Pancreas Tissue ATP in Animal and Preclinical Human Models, J. Clin. Med. 8 (2019).

32. P. M. Porrett, B. J. Orandi, V. Kumar, J. Houp, D. Anderson, A. Cozette Killian, V. Hauptfeld-Dolejsek, D. E. Martin, S. Macedon, N. Budd, K. L. Stegner, A. Dandro, M. Kokkinaki, K. V. Kuravi, R. D. Reed, H. Fatima, J. T. Killian, G. Baker, J. Perry, E. D. Wright, M. D. Cheung, E. N. Erman, K. Kraebber, T. Gamblin, L. Guy, J. F. George, D. Ayares, J. E. Locke, First clinical-grade porcine kidney xenotransplant using a human decedent model, Am. J. Transplant. 00, 1–17 (2022).

33. D. Hoeksma, R. A. Rebolledo, M. Hottenrott, Y. S. Bodar, J. J. Wiersema-Buist, H. Van Goor, H. G. D. Leuvenink, Inadequate Antioxidative Responses in Kidneys of Brain-Dead Rats (2017).

34. E. R. Pieter Hoogland, E. E. De Vries, M. H. L. Christiaans, B. Winkens, M. G. J. Snoeijs, L. W. Ernest Van Heurn, The value of machine perfusion biomarker concentration in DCD kidney transplantations, Transplantation 95, 603–610 (2013).

35. P. Mahboub, P. Ottens, M. Seelen, N. t Hart, H. Van Goor, R. Ploeg, P. Martins, H. Leuvenink, P. Martins, H. Leuvenink, G. Camussi, Ed. Gradual Rewarming with Gradual Increase in Pressure during Machine Perfusion after Cold Static Preservation Reduces Kidney Ischemia Reperfusion Injury, PLoS One 10, e0143859 (2015).

